# Tuning intrinsic disorder predictors for virus proteins

**DOI:** 10.1101/2020.10.27.357954

**Authors:** Gal Almog, Abayomi S Olabode, Art FY Poon

## Abstract

Many virus-encoded proteins have intrinsically disordered regions that lack a stable folded threedimensional structure. These disordered proteins often play important functional roles in virus replication, such as down-regulating host defense mechanisms. With the widespread availability of next-generation sequencing, the number of new virus genomes with predicted open reading frames is rapidly outpacing our capacity for directly characterizing protein structures through crystallography. Hence, computational methods for structural prediction play an important role. A large number of predictors focus on the problem of classifying residues into ordered and disordered regions, and these methods tend to be validated on a diverse training set of proteins from eukaryotes, prokaryotes and viruses. In this study, we investigate whether some predictors outperform others in the context of virus proteins. We evaluate the prediction accuracy of 21 methods, many of which are only available as web applications, on a curated set of 126 proteins encoded by viruses. Furthermore, we apply a random forest classifier to these predictor outputs. Based on cross-validation experiments, this ensemble approach confers a substantial improvement in accuracy, *e.g*., a mean 36% gain in Matthews correlation coefficient. Lastly, we apply the random forest predictor to SARS-CoV-2 ORF6, an accessory gene that encodes a short (61 AA) and moderately disordered protein that inhibits the host innate immune response.

## Introduction

For almost a century, it was assumed that proteins required a properly folded and stable threedimensional or tertiary structure in order to function [1–3]. More recently, it has become evident that many proteins and protein regions are disordered, which are referred to as intrinsically disordered proteins (IDPs) and intrinsically disordered protein regions (IDPRs), respectively. Both IDPs and IDPRs can perform important biological functions despite lacking a properly folded and stable tertiary structure [2, 4].

These kinds of proteins are an important area of research because they play major roles in cell regulation, signalling, differentiation, survival, apoptosis and proliferation [5, 6]. Some are also postulated to be involved in disease etiology and could represent potential targets for new drugs [7, 8]. Virus-encoded IDPs facilitate multiple functions such as adaptation to new or dynamic host environments, modulating host gene expression to promote virus replication, or counteracting host-defense mechanisms [9–11]. IDPRs may be more tolerant of non-synonymous mutations than ordered protein regions [12], which may partly explain why virus genomes can tolerate high mutation rates [13, 14]. Viruses also have very compact genomes with overlapping reading frames [15, 16], in which mutations may potentially modify multiple proteins. This may confer viruses a greater capacity to acquire novel functions and interactions [17]. Overlapping regions tend to be more structurally disordered when compared to non-overlapping regions [18].

Several experimental techniques are available to detect IDPs and IDPRs. The most common methods identify either protein regions in crystal structures that have unresolvable coordinates (X-ray crystallography) or regions in nuclear magnetic resonance (NMR) structures that have divergent structural conformations [6, 19, 20]. Other experimental techniques include circular dichroism (CD) spectroscopy and limited proteolysis (LiP) [1]. The challenge, however, is that these methods are very labour-intensive and difficult to scale up to track the rapidly accumulating number of unique protein sequences in public databases [6, 19]. At the time of this writing, over 60 million protein sequences have been deposited in the Uniprot database, yet only 0.02% of these sequences have been annotated for disorder [6]. As a result, numerous computational techniques that could potentially predict intrinsic disorder in protein sequences have been developed. These techniques work based on the assumptions that compared to IDPs and IDPRs, ordered proteins have a different amino acid composition as well as levels of sequence conservation [21, 22]. To date about 60 predictors for intrinsic disorder in proteins have been developed [1, 23, 24], which can be broadly classified into three major categories. The first category, the scoring function-based methods, predict protein disorder solely based on basic statistics of amino acid propensities, physio-chemical properties of amino acids and residue contacts in folded proteins to detect regions of high energy. A second category is characterized by the use of machine learning classifiers (e.g., regularized regression models or neural networks) to predict protein disorder based on amino acid sequence properties. The third category are meta-predictors that predict disorder from an ensemble of predictive methods from the other two categories [1, 6, 25].

Different predictors of intrinsic disorder are developed on a variety of methodologies and will inevitably vary with respect to their sensitivities and biases in application to different protein sequences. As a result, it has been relatively difficult to benchmark these methods to identify a single disorder prediction method that can be classified as the most accurate relative to the others [24]. The DisProt database is a good resource for obtaining experimental data that has been manually curated for disorder in proteins, and can be used for benchmarking the performance of disorder predictors. As of April 27th, 2020, the Disprot protein database contained *n* = 3500 proteins of which 126 were virus-encoded proteins that have been annotated for intrinsic disorder as a presence-absence characteristic at the amino acid level [26, 27]. Previously, Tokuriki *et al*. [14] reported preliminary evidence that when compared to non-viruses, viral proteins possess many distinct biophysical properties including having shorter disordered regions. We are not aware of a published study that has previously benchmarked predictors of intrinsic disorder specifically for viral proteins. Here, we report results from a comparison of 21 disorder predictors on viral proteins from the DisProt database to firstly determine which methods work best for viruses, and secondly to generate inputs for an ensemble predictor that we evaluate alongside the predictors used individually.

## Methods

### Data collection

The Database of Protein Disorder (DisProt) [28] was used to collect virus protein sequences annotated with intrinsically disordered regions, based on experimental data derived from various detection methods; *e.g*., X-ray crystallography, NMR spectroscopy, CD spectroscopy (both far and near UV) and protease sensitivity. DisProt records include the amino acid sequence and all disordered regions annotated with the respective detection methods as well as specific experimental conditions. At the time of our study, DisProt contained 3,500 author-verified proteins, of which all viral proteins were collected for the present study. A total of 126 virus proteins were obtained, derived from different detection methods. Similarly, a set of 126 non-viral proteins was sampled at random without replacement from the protein database for comparison.

We evaluated a number of disorder prediction programs and web applications. From the methods tested, we selected a subset of predictors favouring those that were developed more recently, are actively maintained, and performed well in previous method comparison studies [1, 29]. Where alternate settings or different versions based on training data were available for a given predictor, we tested all combinations. Our final set of 21 prediction methods tested were: SPOT-Disorder2 [30], PONDR-FIT [31], IUPred2 (short and long) [32], PONDR (VLXT, XL1-XT, CAN-XT, VL3-BA, and VSL2 variants) [33], Disprot (VL2 and variants VL2-V, -C and -S; VL3, VL3H, and VSLB) [34], CSpritz (short and long) [35], and ESpritz (variants trained on X-ray, NMR, and Disprot data) [36]. Although several other predictor models have been released online, the respective web services were unavailable or broken over the course of our data collection.

To obtain disorder predictions from the methods that were only accessible as web applications, *i.e*., with no source code or compiled binary standalone distribution, we wrote Python scripts to automate the process of submitting protein sequence inputs and parsing HTML outputs. We used Selenium in conjunction with ChromeDriver (v81.0.4044.69) [37] to automate the web browsing and form submission processes. For each predictor, we implemented a delay of 90 seconds between consecutive protein sequence queries to avoid overloading the webservers hosting the respective predictor algorithms with repeated requests. Due to issues with the Disprot webserver, we were only able to obtain predictions for the non-viral protein data set for 13 of the predictors.

We converted each DisProt record to a binary vector corresponding to ordered/disordered state of residues in the amino acid sequence. To compare results between disorder prediction algorithms, we dichotomized continuous-valued residue predictions, i.e., intrinsic disorder probability, by locating the threshold that maximized the Matthews correlation coefficient (MCC) for each predictor applied to the DisProt training data. This optimal threshold was estimated using Brent’s root-finding algorithm as implemented by the *optim* function in the R statistical computing environment (version 3.4.4). In addition, we calculated the accuracy, specificity and sensitivity for each predictor from the contingency table of DisProt residue labels and dichotomized predictions.

### Ensemble classifier training and validation

To assess whether the accuracy of existing predictors could be further improved on the virus-specific data set, we trained an ensemble classifier on the outputs of all predictors as features. Specifically, we used the random forest method implemented in the *scikit-learn* (version 0.23.1) Python module [38], which employs a set of de-correlated decision trees and averages their respective outputs to obtain an ensemble prediction [39]. To reduce bias, random forests fit the same decision trees to many bootstrap samples of the training data, and each committee of trees ‘votes’ for a particular classification [40]. By splitting the trees based on different samples of features, random forests reduce the correlation between trees and the overall variance.

We split the viral protein data into random testing and training subsets, with 30% of protein sequences reserved for testing. Due to class imbalance in the data (*i.e*., only a minority of residues are labeled as disordered), we used stratified random sampling using the ‘StratifiedShuffleSplit’ function in the *scikit-learn* module. This function stratifies the data by label so that a constant proportion of labels is maintained in the training subset. Continuous-valued outputs from each predictor were normalized to a zero mean and unit variance. Thus, we did not apply the dichotomizing thresholds to these features (predictor outputs) when training the random forest classifier.

We used 5-fold cross validation to tune the four hyper-parameters of the random forest classifier; namely: (1) the number of decision trees; (2) the maximum depth of any given decision tree; and the minimum number of samples required to split (3) an internal node or (4) a leaf node. To further minimize the effect of class imbalance in our data, we used over-sampling to balance the data with synthetic cases [41]. As suggested in [42], we applied an over-sampling procedure at every iteration of the cross-validation analysis to avoid over-optimistic results. We used the Python package *imbalanced-learn* [43] to over-sample the minority class (residues in intrinsically disordered regions) using the synthetic minority oversampling technique (SMOTE) [44]. SMOTE generates new cases by sampling the original data at random with replacement, evaluates each sample’s *k* nearest neighbours in the feature space, and then generates new synthetic samples along the vectors joining the sample to one of the neighbouring points. Over-sampling enables decision trees to be more generalizable by amplifying the decision region of the minority class.

Using the optimized tuning parameters, we fit the final model on all of the training data. We applied this final model to generate predictions on the reserved testing data and calculated the MCC, sensitivity, specificity and accuracy. We repeated this process 10 times with randomly generated seeds to split the data into training and testing subsets, and averaged these performance metrics across replicates.

### Comparison to non-viral data

To characterize how the performance of individual disorder predictors might vary among proteins from viruses and non-viruses, we computed the root mean square error (RMSE) for all continuous-valued predictions relative to the Disprot label (0, 1). We visualized this error distribution using principal component analysis (PCA). As well, we trained a support vector machine (SVM) on the RMSE values to determine whether the virus/non-virus labels were separable in this space. We used the default radial basis kernel with the C-classification SVM method implemented in the R package *e1071* [45], with 100 training subsets sampled at random without replacement for half of the data, and the remaining half for validation.

### Data availability

We have released the Python scripts for automating queries to the disorder prediction web servers under a permissive free software license at https://github.com/PoonLab/Floppy/.

## Results and Discussion

### Viral and non-viral proteins have similar levels of disorder

We obtained 126 viral and 126 randomly selected non-viral protein sequences from the DisProt database. The sequences were already annotated manually by a panel of experts for the presence or absence of disorder at each amino acid position, based on experimental data [27]. Supplementary Tables S1 and S2 summarize the composition of the viral and non-viral protein datasets, respectively. The viral protein data set represents 22 virus families and 48 species. Not surprisingly, human immunodeficiency virus type 1 was disproportionately represented in these data with 16 entries corresponding to seven different gene products. Similarly, the non-viral protein data set was predominated by 75 human proteins, followed by 23 proteins from the yeast *Saccharomyces cerevisiae*. We found no significant difference in amino acid sequence lengths between viruses and all other organisms (Wilcoxon rank sum test, *P* = 0.60), with median lengths of 355 [interquartile range, IQR: 145–846] and 395 [203–729] amino acids, respectively. Furthermore, the dispersion in sequence lengths was significantly greater among viral proteins relative to the nonviral proteins (Ansari-Bradley test, *P* = 0.0028). There was no significant difference in the proportion of residues in disordered regions between the viral and non-viral data (Wilcoxon *P* = 0.97). The mean proportions were 0.30 (interquartile range, IQR [0.07-0.42]) for viral and 0.30 [0.07-0.47] for non-viral proteins, and similar numbers of proteins exhibited complete disorder (13 and 9, respectively).

### Divergent predictions of disorder in viral proteins

Our first objective was to benchmark the performance of different predictors of intrinsic protein disorder to determine which predictor conferred the highest accuracy for viral proteins. These predictors generate continuous-valued outputs that generally correspond to the estimated probability that the residue is in an intrinsically disordered region. To create a uniform standard for comparison to the binary presence-absence labels, we optimized the disorder prediction thresholds as a tuning parameter for each predictor for the viral and non-viral datasets, respectively (Supplementary Tables S3 and S4). Put simply, residues with values above the threshold were classified as disordered. We used both the Matthews correlation coefficient (MCC, ranging from −1 to +1 [46]) and area under the receiver-operator characteristic curve (AUC, ranging from 0 to 1) to quantify the performance of each predictor.

These quantities were significantly correlated (Spearman’s *ρ* = 0.95, *P* = 5.2 × 10^−6^) and identified ESpritz.Disprot, CSpritz.Long and SPOT.Disorder2 as the most effective predictors for the viral proteins (Figure 1A). ESpritz.Disprot obtained the highest overall values for both MCC and AUC (0.46 and 0.85, respectively). We note that SPOT.Disorder2 has recently been reported to exhibit a high degree of prediction accuracy for proteins of varying length [47]. In contrast, the predictors Disprot-VL2-V, PONDR-XL1 and PONDR-CAN performed very poorly on the viral dataset with MCC < 0.2 and AUC < 0.65. VL2-V is a ‘flavour’ of the VL2 predictors which were allowed to specialize on different subsets of a partitioned training set; for example, V tended to call higher levels of disorder in proteins of Archaebacteria [34]. Similarly, PONDR-XL1 was optimized to predict longer disordered regions and PONDR-CAN was trained specifically on calcineurins (a protein phosphatase) that is known to perform poorly on other proteins [48].

**Figure 1:**
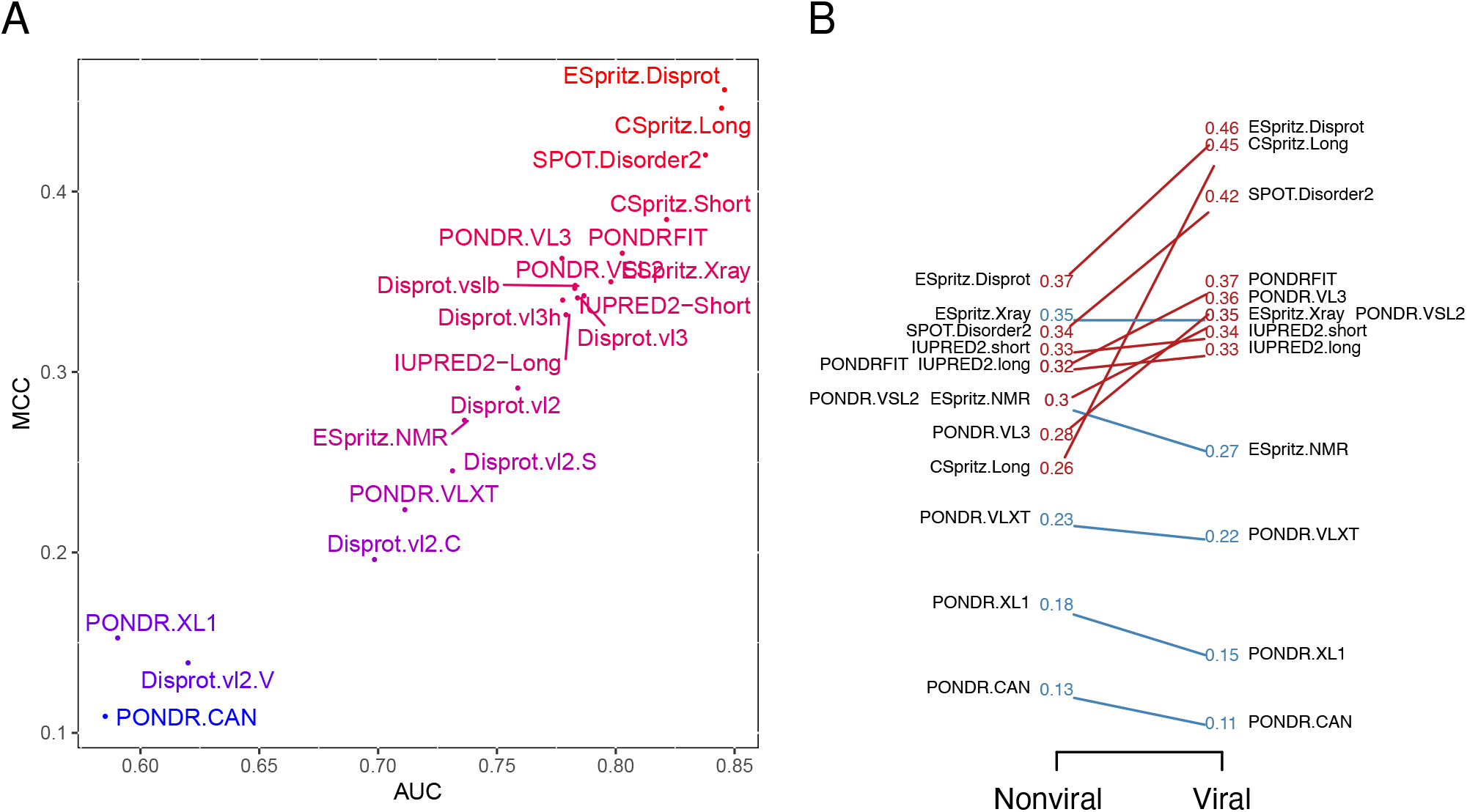
Performance of predictors on viral data set. (A) Scatterplot of MCC and AUC values for 21 predictors applied to the viral protein data set. (B) Slopegraph comparing the MCC values for 13 predictors applied to both non-viral and viral data sets. Because the three variants of the ESpritz model obtained identical MCC values, the corresponding labels were merged. Two labels (PONDRFIT, PONDR.VSL2) were displaced to prevent overlaps on the left and right sides, respectively.

Figure 1B compares the MCC values for non-viral and viral protein data sets. Predictors exhibited substantially less variation in MCC for the non-viral data — put another way, the majority of predictors were more accurate at predicting disorder in viral proteins. The entire set of MCC, AUC, sensitivity and specificity values for both data sets are summarized in Supplementary Tables S3 and S4. To examine potential differences among predictors in greater detail, we calculated the RMSE for each protein and predictor and used a principal components analysis to visualize the resulting matrix (Figure 2). The PCA indicated that the different predictors did not exhibit markedly divergent error profiles at the level of entire proteins. However, a support vector machine classifier trained on a random half of these data obtained, on average, an AUC of 0.75 (n = 100, range = 0.65 − 0.83), indicating that the viral and non-viral protein labels were appreciably separable with respect to these RMSE values.

**Figure 2:**
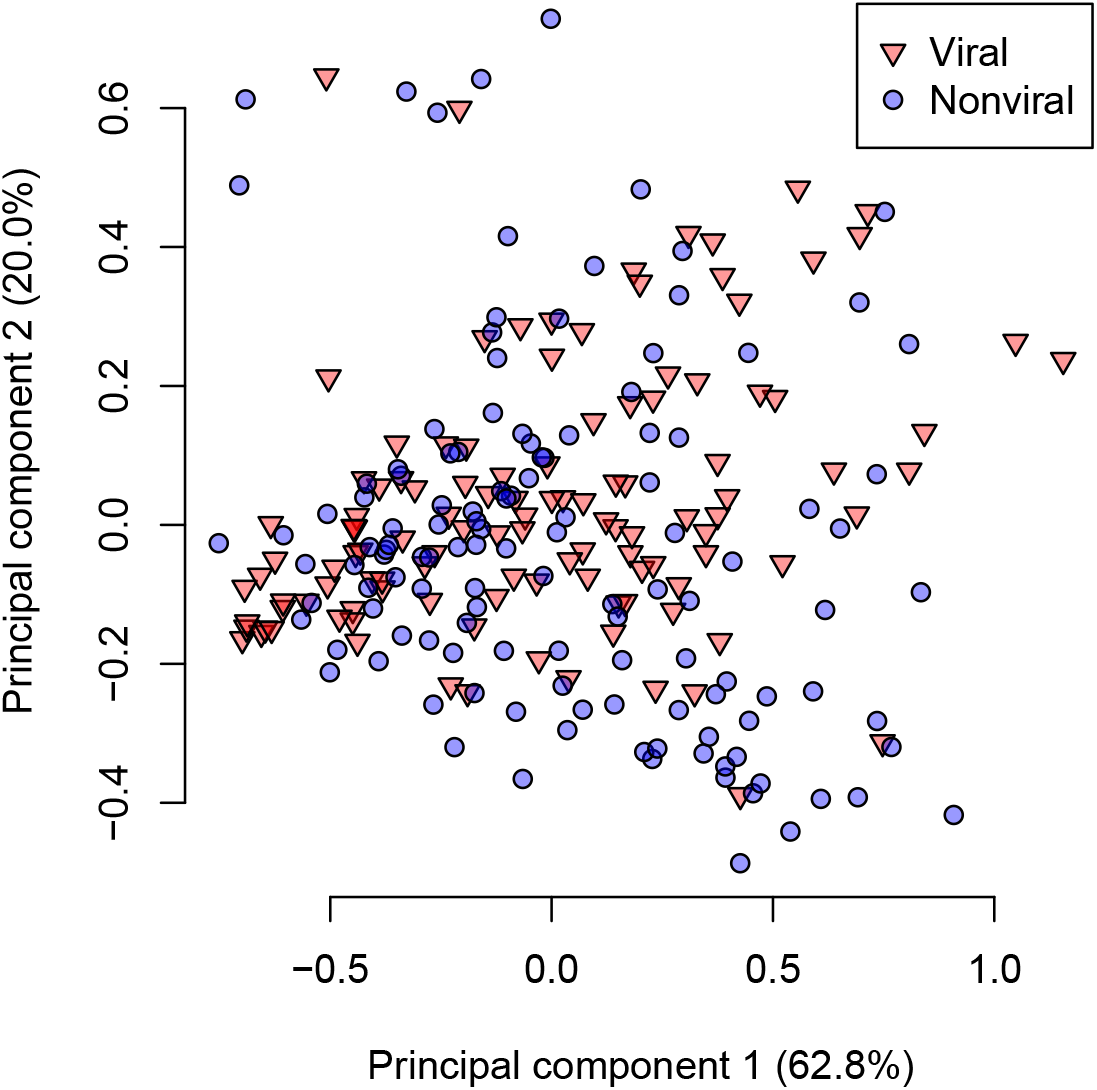
Principal components analysis plot of the root mean squared errors (RMSEs) for 13 disorder predictors on viral (red, triangles) and non-viral (blue, circles) protein sequences. The percentages of total variance explained by the first two principal components are indicated in parentheses in the respective axis labels.

### Ensemble prediction

Ensemble classifiers are expected to perform better than their constituent models because they can reduce overfitting of the data by the latter [49]. Although multiple predictive models of protein disorder employ an ensemble approach, none of them has been trained specifically on viral protein data. We trained a random forest classifier on the outputs of the predictors used in our study using 10 random training subsets of the viral protein data. Next, we validated the performance of this ensemble model in comparison to these individual predictors to determine if training on viral data conferred a significant advantage. We found that the ensemble classifier performed substantially better, with a mean MCC of 0.72 (range 0.62 − 0.86). This corresponded to a roughly 27% improvement relative to ESpritz.Disorder, the best performing disorder predictor on these data (Figure 1).

To examine the relative contribution of the different predictors used as inputs for the ensemble method, we evaluated the feature importance of each input (Figure 3) — roughly the prevalence of that feature among the decision trees comprising the random forest. We observed that the individual accuracy of a predictor did not necessarily correspond to its feature importance. Specifically, the best predictors (ESpritz.Disprot, CSpritz.Long and SPOT-Disorder.2) tended to be assigned higher importance values. On the other hand, both Disprot-VL2.C and Disprot-VL2.V also displayed high importance despite having some of the worst accuracy measures when evaluated individually (Figure 1).

**Figure 3:**
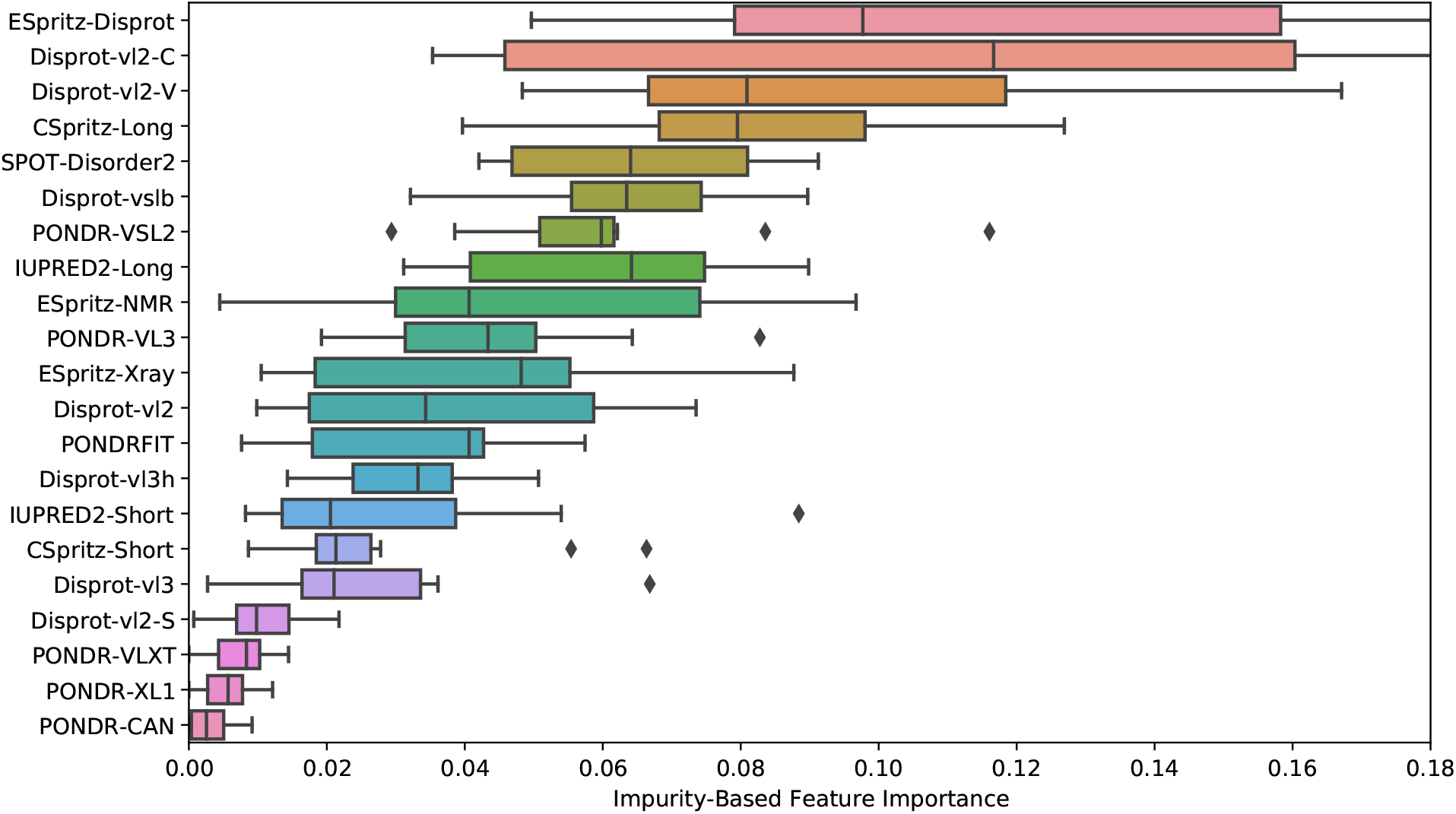
Box plot of the average decrease in Gini impurity by each feature in the random forest, for 10 random runs of the random forest model. Vertical line indicates the median, the box is the interquartile range (IQR; range from first to third quartiles). The left whisker extends to the first datum greater that Q1 − 1.5 × IQR and the right whisker extends to the last datum smaller than Q3 + 1.5 × IQR. Individual points are outliers that lie outside this range.

### Example: SARS-CoV-2 accessory protein 6

To illustrate the use of our ensemble model on a novel protein, we applied this model and the 21 individual predictors to the accessory protein encoded by ORF6 in the novel 2019 coronavirus that was first isolated in Wuhan, China (designated SARS-CoV-2). ORF6 is one of the eight accessory genes of this virus. Its protein product is involved in antagonizing interferon activity thereby suppressing host immune response [50]. The protein is predicted to be highly disordered, particularly in its C-terminal region that contains short linear motifs involved in numerous biological activities [51]. We used a heatmap (Figure 4) to visually summarize results from the ensemble method and individual predictors, mapped to the ORF6 amino acid sequence. Overall, most predictors assigned a higher probability of disorder in the C-terminal region of the protein, with the conspicuous exception of PONDR-XL1 and PONDR-CAN, which did not predict any disordered residues in this region. We also observed considerable variation among predictors around this overall trend. Although the PONDR-XL1 predictor is documented to omit the first and last 15 residues from disorder predictions, we observed that only 14 residues were reported this way — this treatment was also obtained for PONDR-CAN, although it was not a documented behaviour of that predictor.

**Figure 4:**
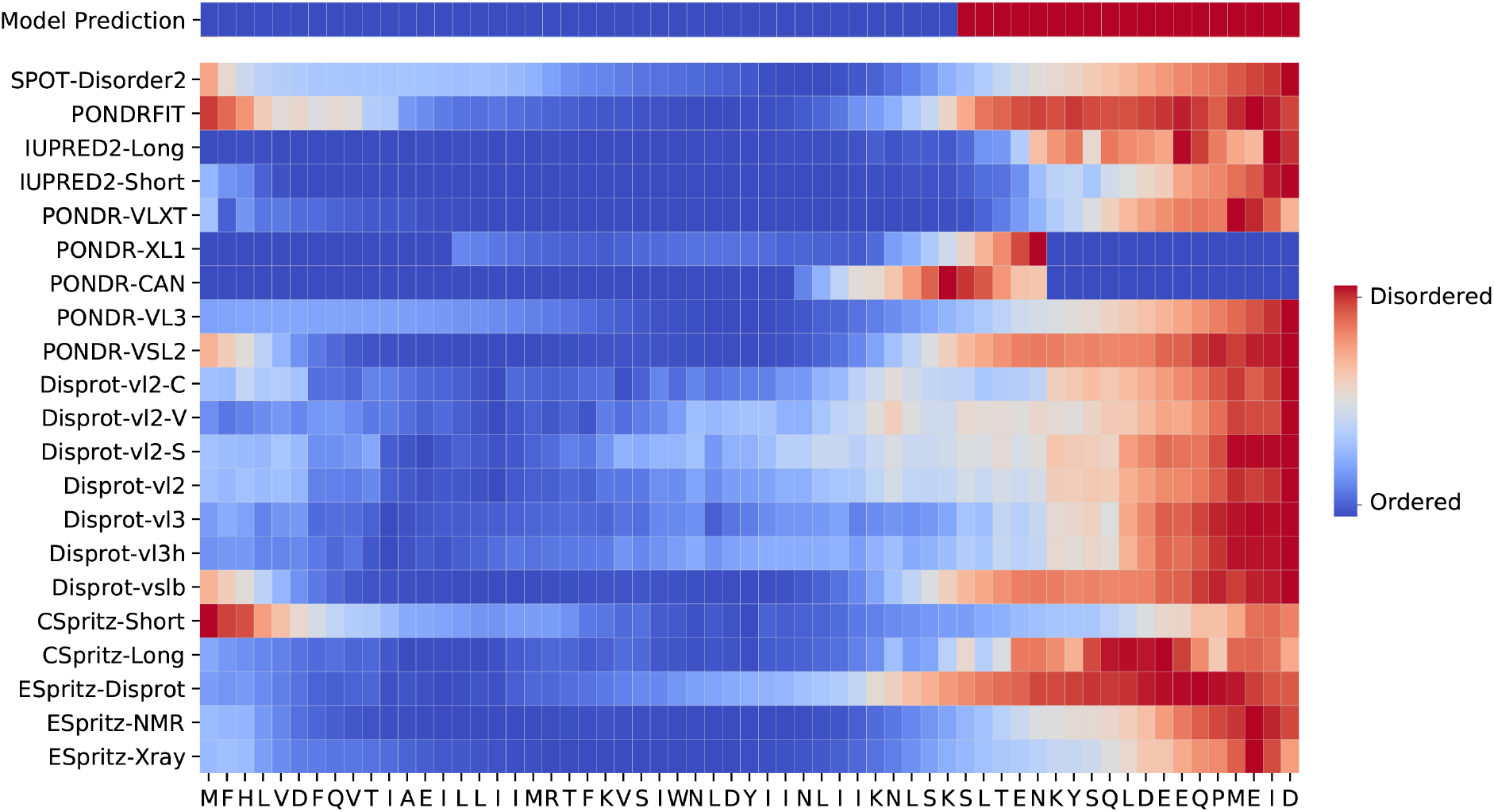
Disorder Predictions for novel ORF6 in SARS-CoV-2. The first row represents the random forest model predictions, with subsequent rows corresponding to individual predictors. The entire protein length is represented on the x-axis, each grid is an amino acid. Red squares indicate disordered predictions and blue squares indicate ordered predictions.

## Concluding remarks

Intrinsically disordered protein regions play an essential role in many viral functions [11]. It is therefore important to predict these regions accurately in order to make biological inferences from sequence variation. In this study, we found that predictive models of intrinsic disorder were more divergent in performance when evaluated on viral proteins than non-viral proteins. We note that many of these predictors could only be accessed through web applications, and some services become unavailable at different points of our study. Although we obtained more accurate predictions — or at least, predictions that were more concordant with an expert-curated database of intrinsic protein disorder [27] — using an ensemble ‘machine learning’ method, the erratic availability of the constituent predictors presents a significant obstacle to the practical utility of such approaches. Hence, we encourage researchers in the field of disorder prediction to support open science by releasing their source code or compiled binaries for local execution.

## Acknowledgments

This work was supported in part by grants from the Natural Sciences and Engineering Research Council of Canada (NSERC, RGPIN-2018-05516) and from the Canadian Institutes of Health Research (CIHR, PJT-155990).

## Supplementary Tables

**Table S1:**
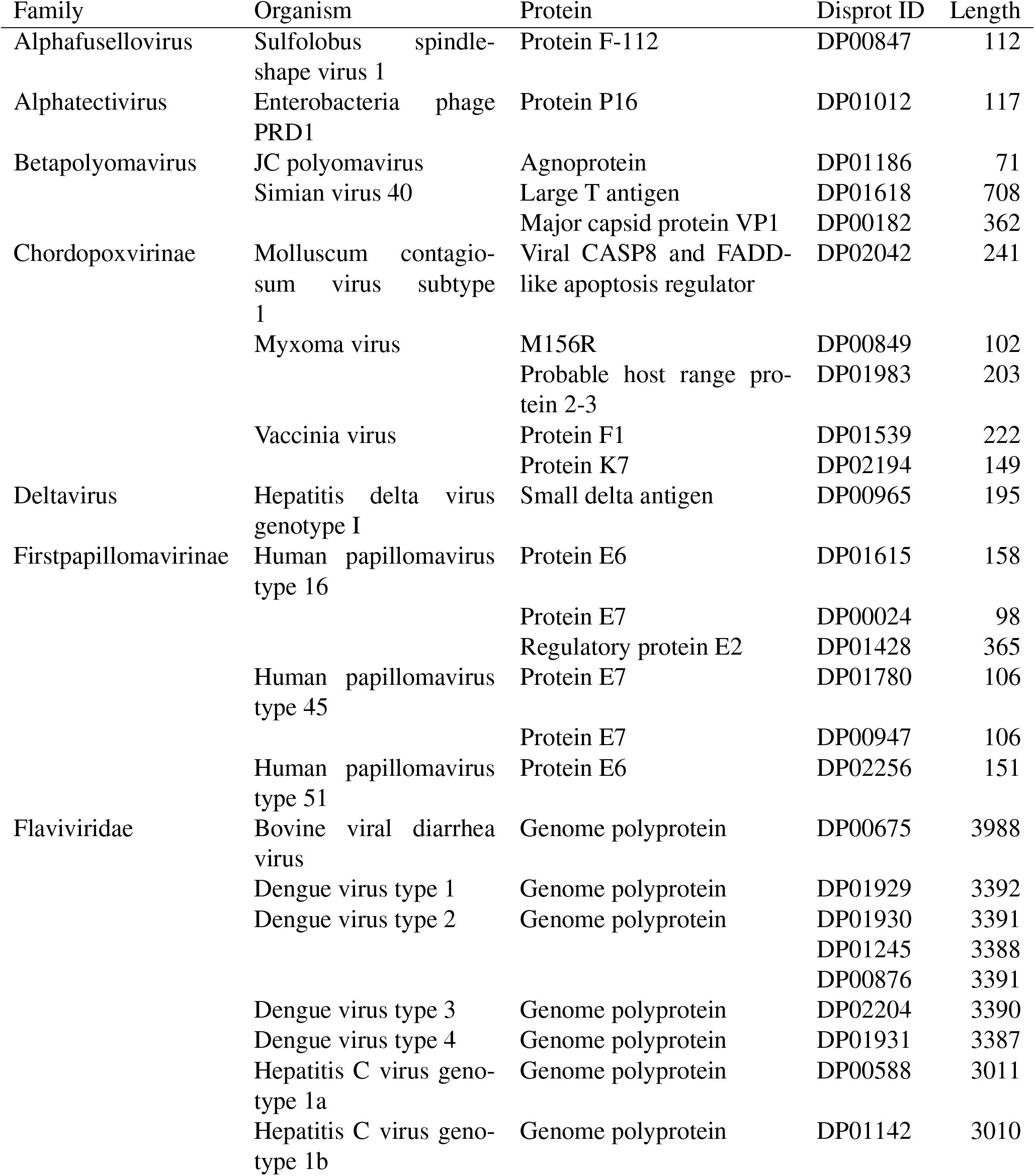
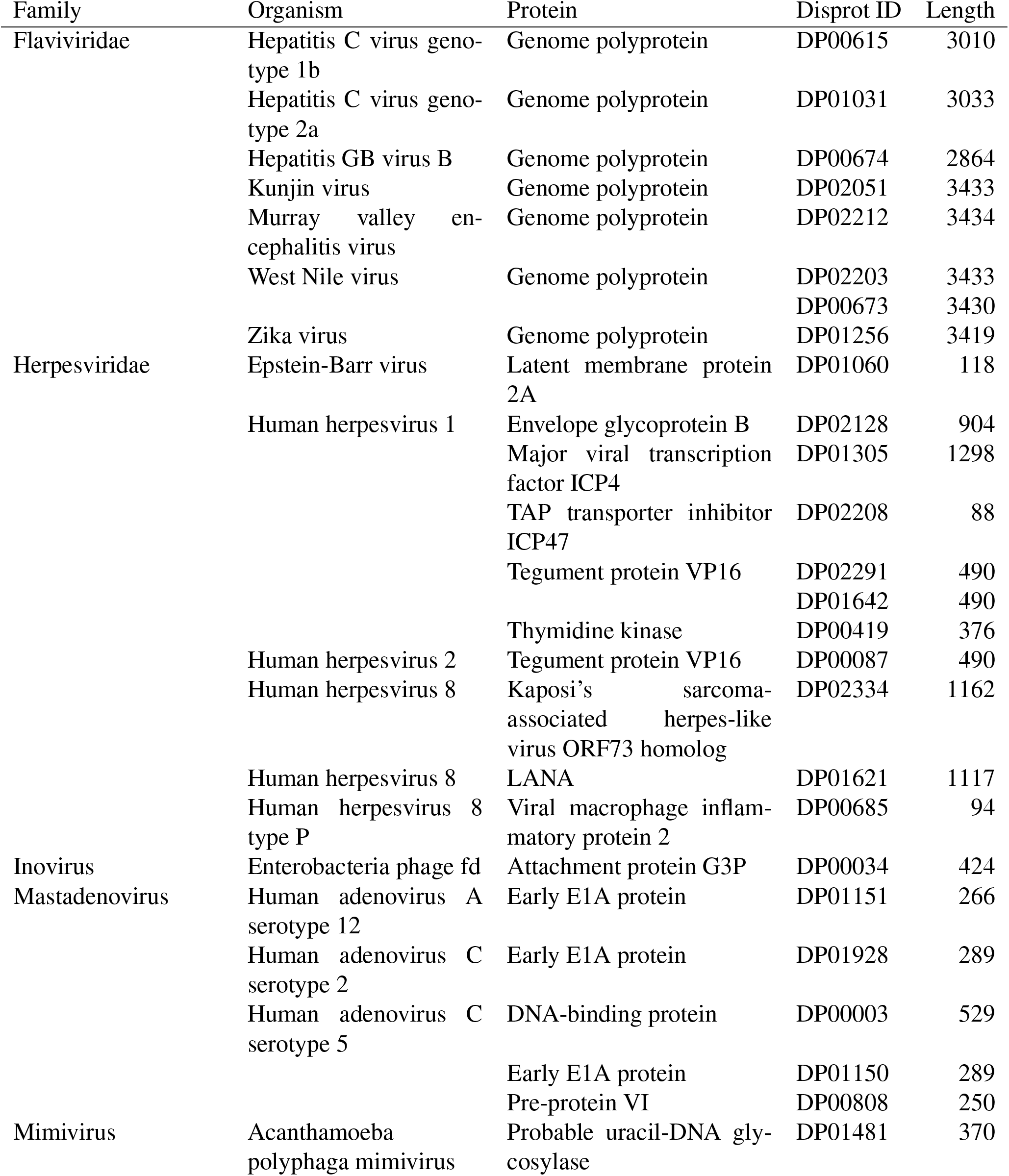
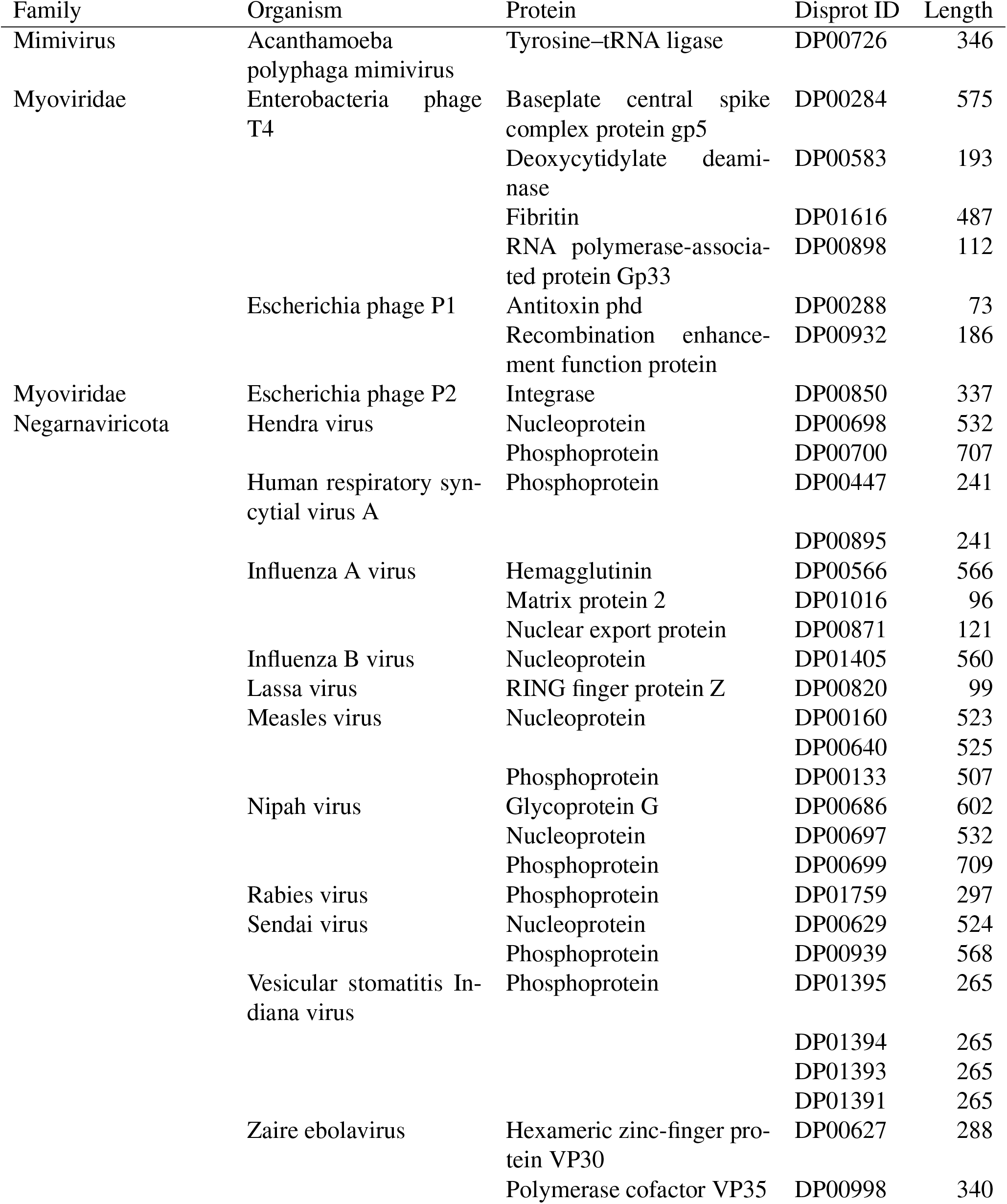
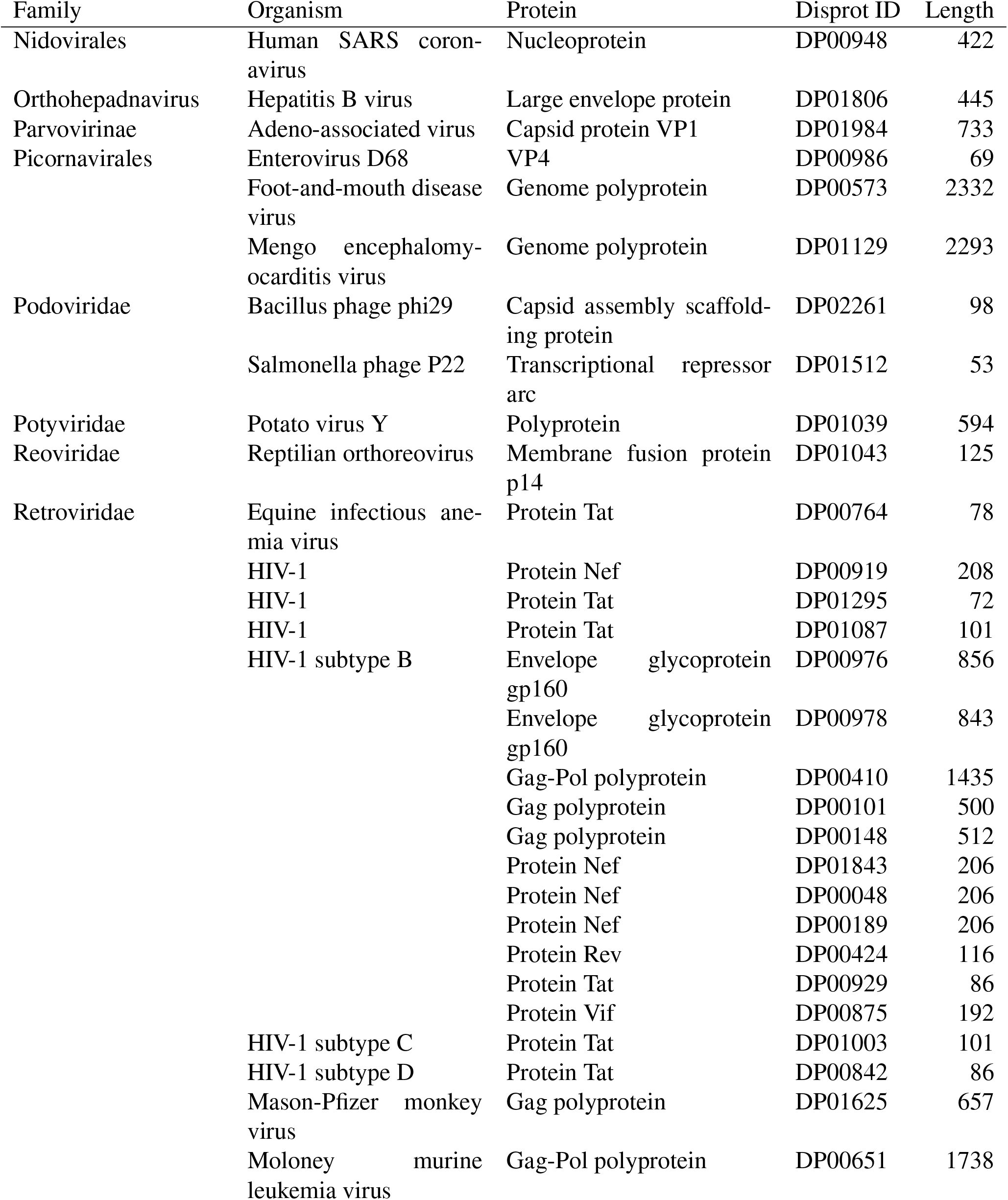
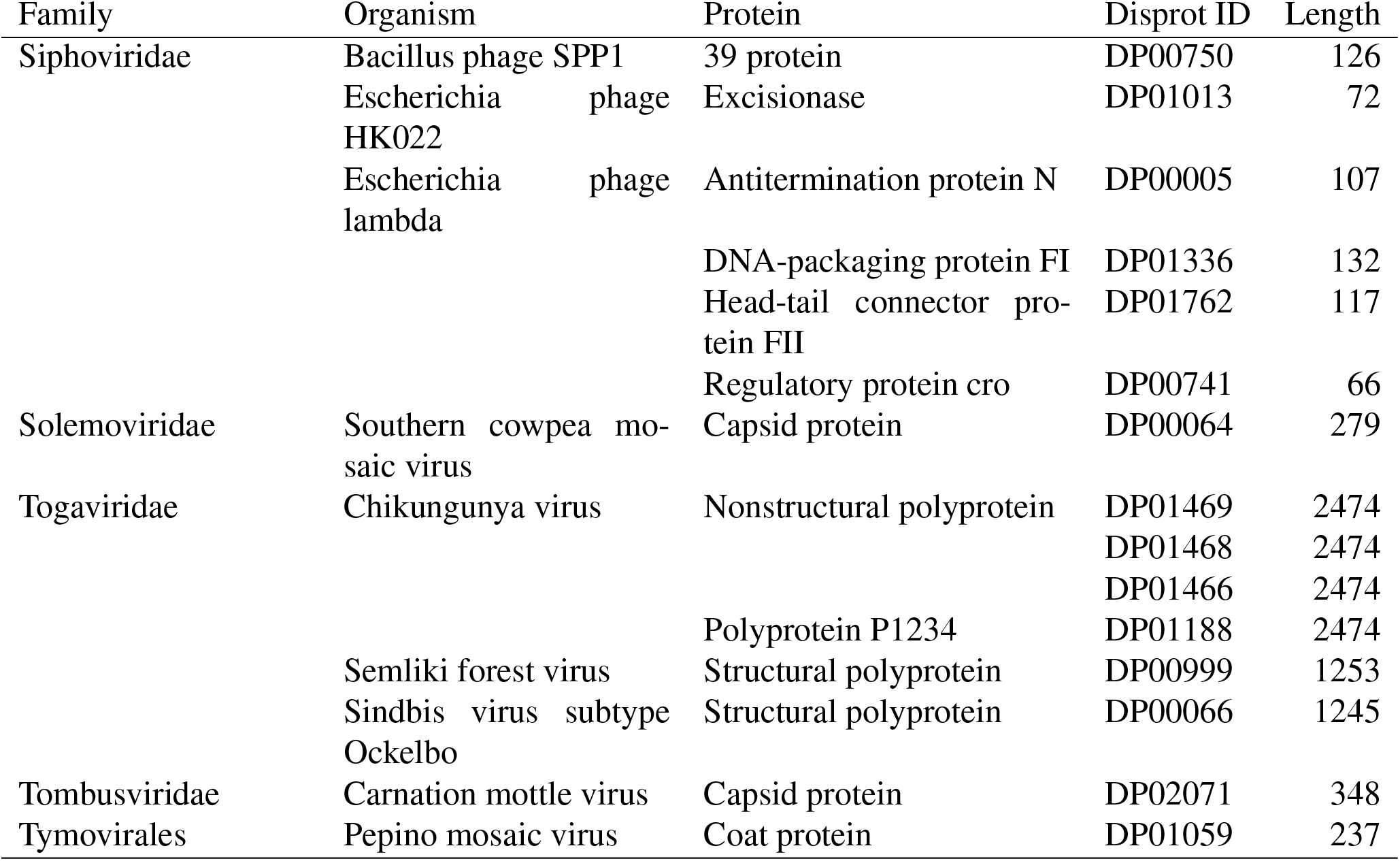
Summary of viral protein data

**Table S2:**
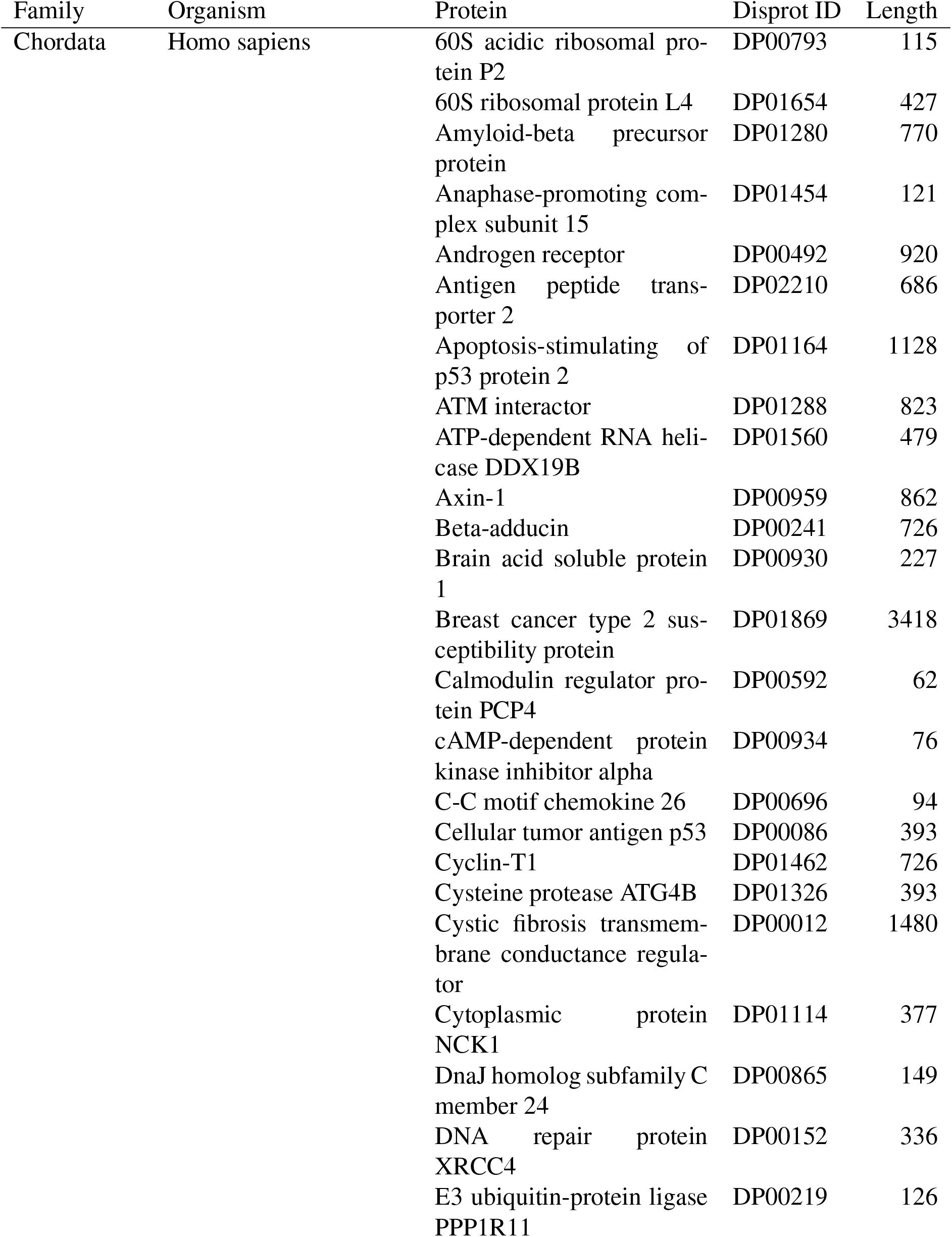
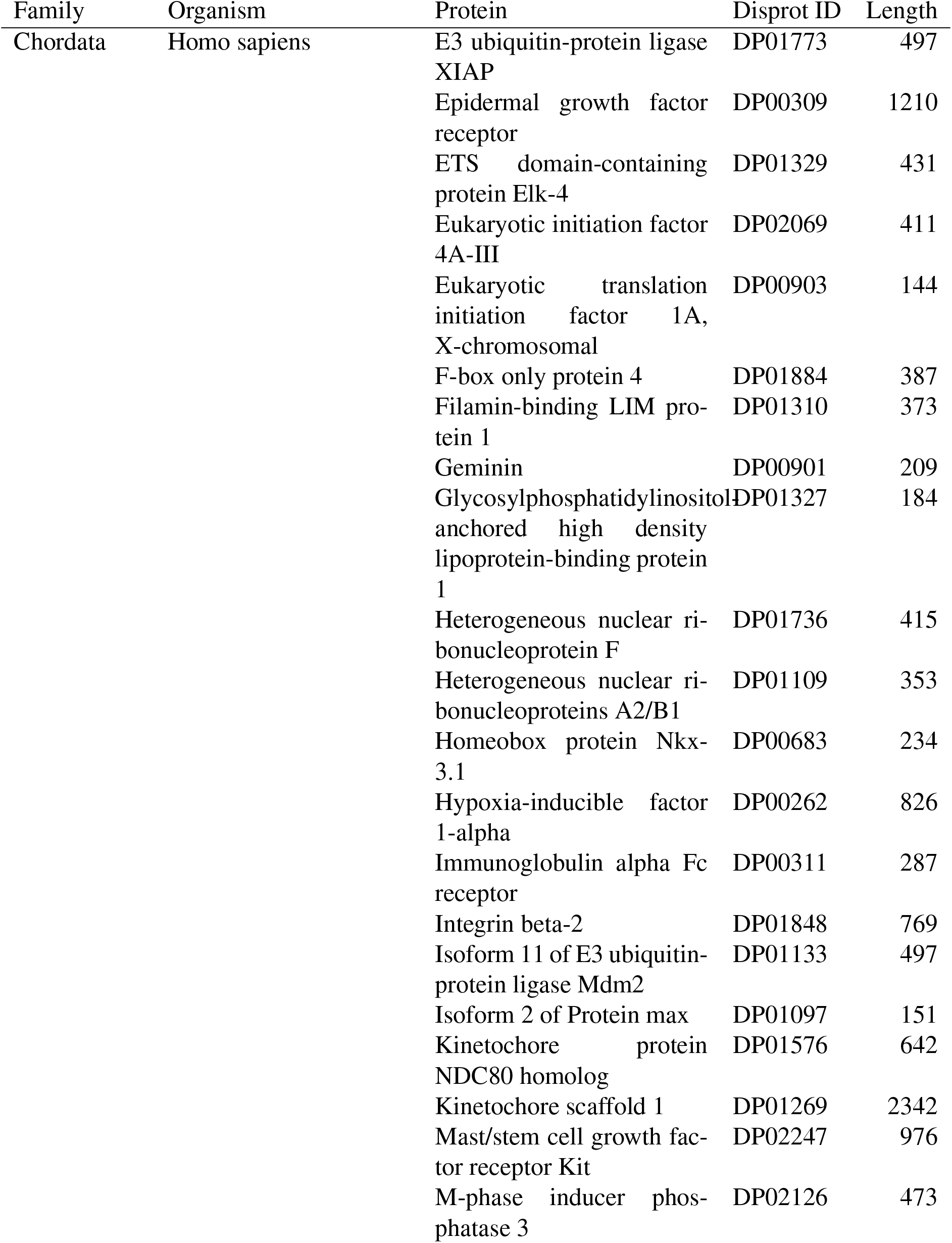
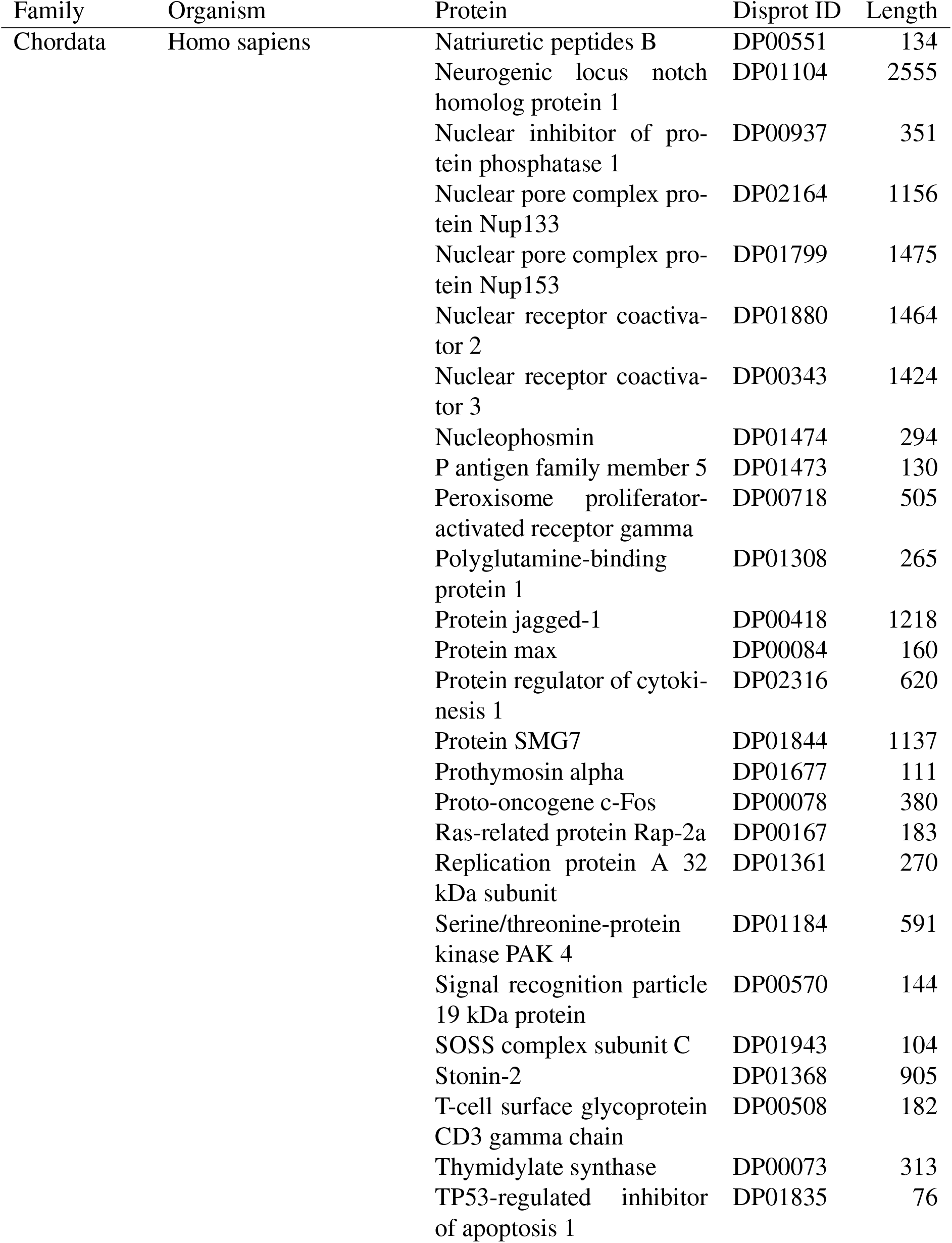
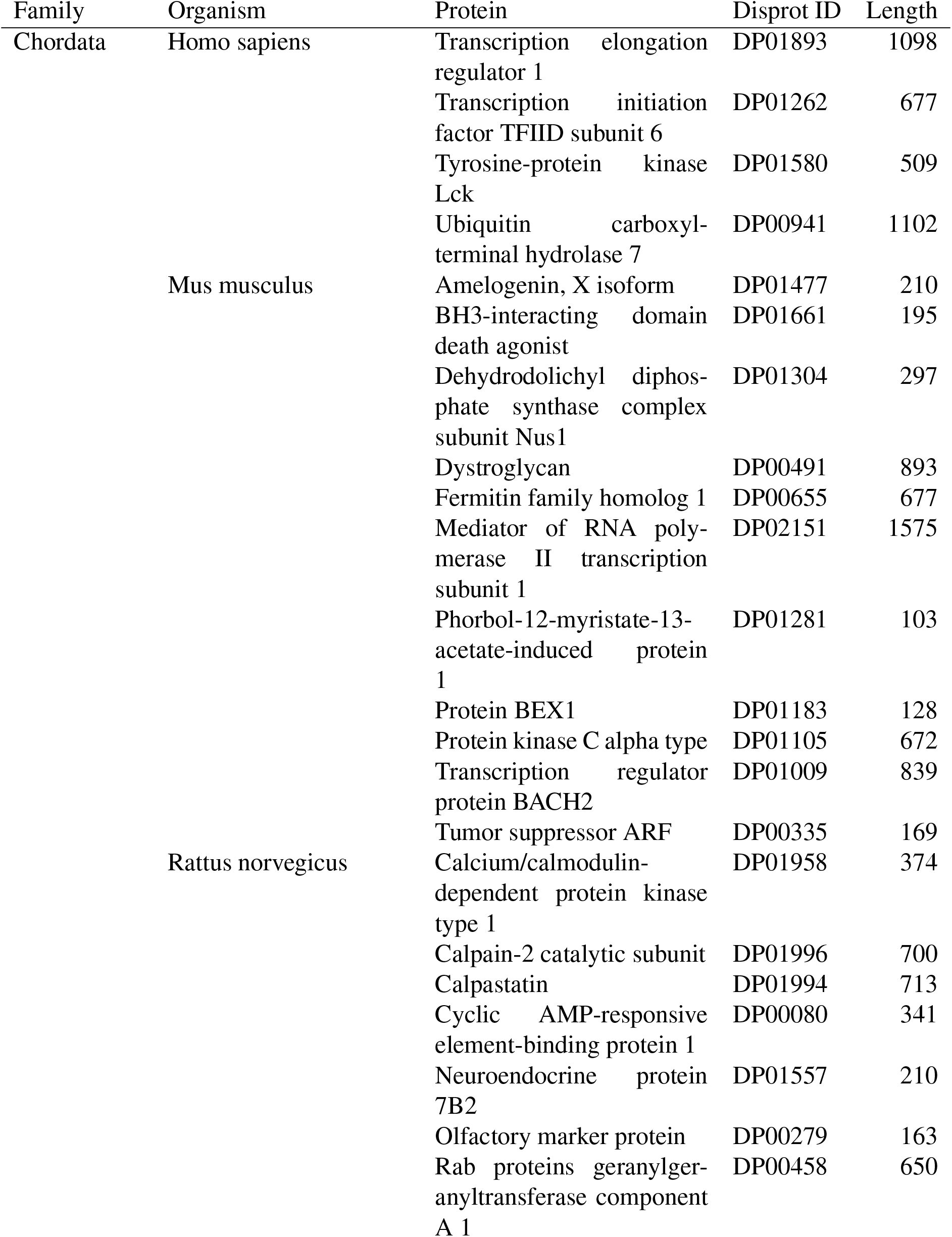
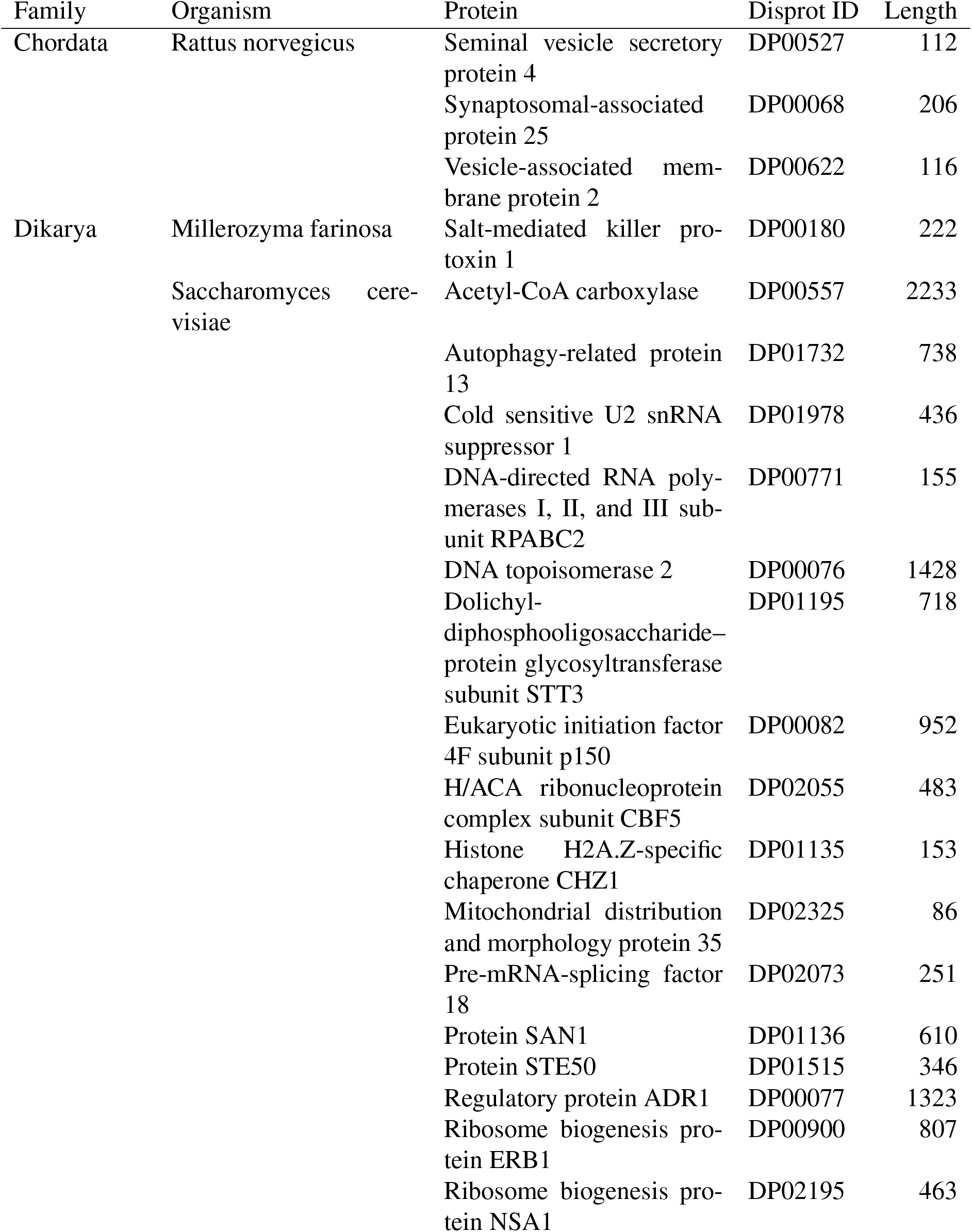
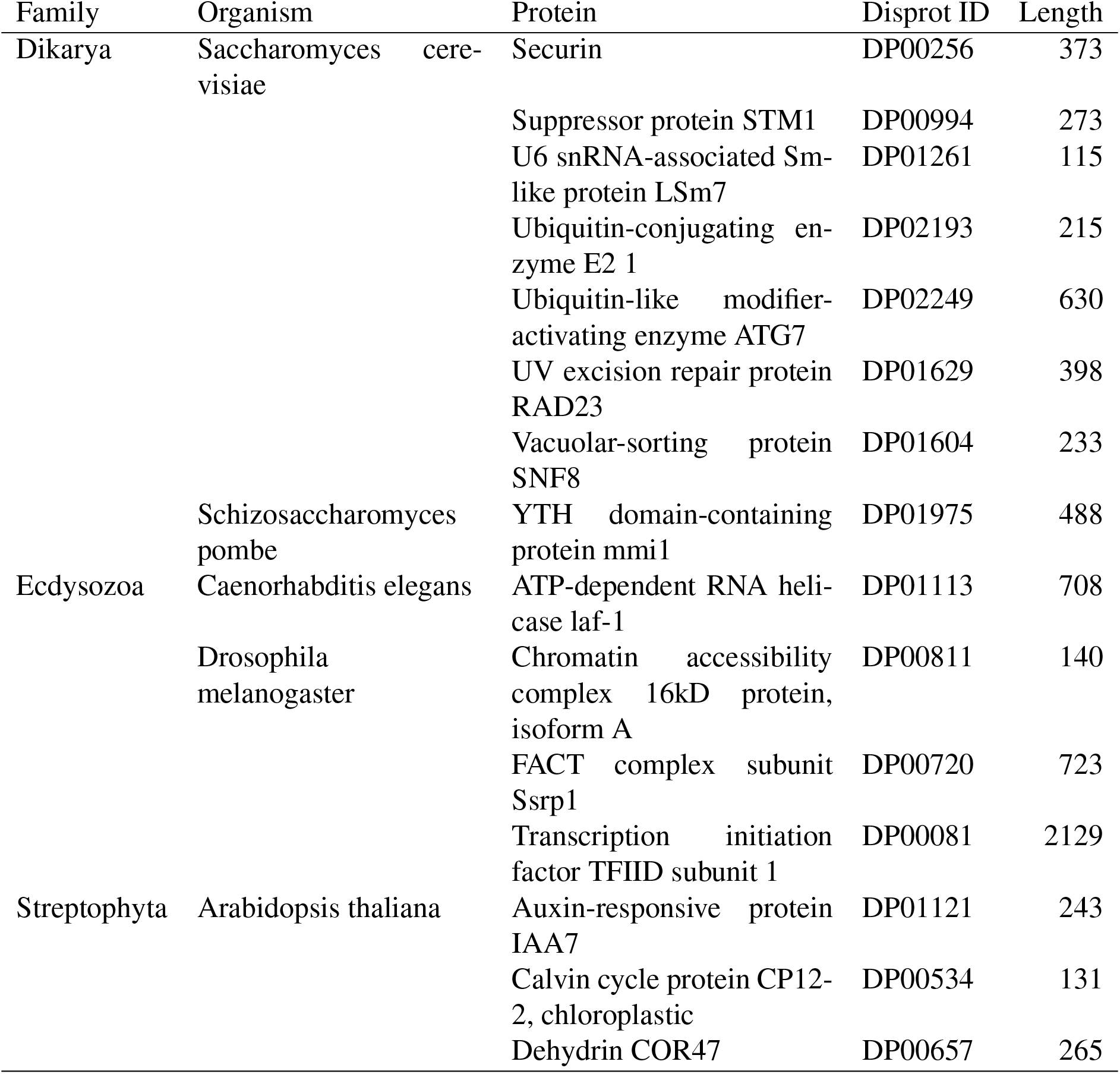
Summary of non-viral protein data

**Table S3:** Optimized thresholds and accuracy of intrinsic disorder prediction in viral proteins for each predictor analysed. MCC = Matthews correlation coefficient; AUC = area under the receiver operator characteristic curve.

| Predictor | MCC | AUC | Specificity | Sensitivity | Threshold |
| --- | --- | --- | --- | --- | --- |
| ESpritz.Disprot | 0.46 | 0.85 | 0.64 | 0.90 | 0.29 |
| CSpritz.Long | 0.45 | 0.85 | 0.57 | 0.92 | 0.34 |
| SPOT.Disorder2 | 0.42 | 0.84 | 0.65 | 0.88 | 0.13 |
| CSpritz.Short | 0.38 | 0.82 | 0.67 | 0.85 | 0.11 |
| PONDRFIT | 0.37 | 0.80 | 0.57 | 0.88 | 0.50 |
| PONDR.VL3 | 0.36 | 0.78 | 0.56 | 0.88 | 0.54 |
| ESpritz.Xray | 0.35 | 0.80 | 0.58 | 0.87 | 0.06 |
| Disprot.vslb | 0.35 | 0.78 | 0.54 | 0.88 | 0.65 |
| PONDR.VSL2 | 0.35 | 0.78 | 0.54 | 0.88 | 0.65 |
| IUPRED2.short | 0.34 | 0.79 | 0.57 | 0.87 | 0.47 |
| Disprot.vl3 | 0.34 | 0.79 | 0.53 | 0.88 | 0.68 |
| Disprot.vl3h | 0.34 | 0.78 | 0.55 | 0.87 | 0.63 |
| IUPRED2.long | 0.33 | 0.78 | 0.54 | 0.87 | 0.49 |
| Disprot.vl2 | 0.29 | 0.76 | 0.53 | 0.85 | 0.57 |
| ESpritz.NMR | 0.27 | 0.74 | 0.52 | 0.84 | 0.36 |
| Disprot.vl2.S | 0.24 | 0.73 | 0.53 | 0.81 | 0.54 |
| PONDR.VLXT | 0.22 | 0.71 | 0.50 | 0.80 | 0.62 |
| Disprot.vl2.C | 0.20 | 0.70 | 0.57 | 0.73 | 0.52 |
| PONDR.XL1 | 0.15 | 0.59 | 0.38 | 0.82 | 0.69 |
| Disprot.vl2.V | 0.14 | 0.62 | 0.29 | 0.87 | 0.52 |
| PONDR.CAN | 0.11 | 0.59 | 0.25 | 0.88 | 0.62 |

**Table S4:** Optimized thresholds and accuracy of intrinsic disorder prediction in non-viral proteins for each predictor analysed. MCC = Matthews correlation coefficient; AUC = area under the receiver operator characteristic curve.

| Predictor | MCC | AUC | Specificity | Sensitivity | Threshold |
| --- | --- | --- | --- | --- | --- |
| ESpritz.Disprot | 0.36 | 0.79 | 0.65 | 0.78 | 0.38 |
| ESpritz.Xray | 0.35 | 0.77 | 0.73 | 0.70 | 0.06 |
| SPOT.Disorder2 | 0.34 | 0.77 | 0.70 | 0.73 | 0.24 |
| IUPRED2.short | 0.33 | 0.76 | 0.72 | 0.69 | 0.42 |
| IUPRED2.long | 0.32 | 0.76 | 0.72 | 0.69 | 0.47 |
| PONDRFIT | 0.32 | 0.75 | 0.74 | 0.67 | 0.46 |
| PONDR.VSL2 | 0.30 | 0.74 | 0.76 | 0.63 | 0.62 |
| PONDR.VL3 | 0.28 | 0.74 | 0.78 | 0.59 | 0.51 |
| ESpritz.NMR | 0.28 | 0.73 | 0.65 | 0.70 | 0.39 |
| CSpritz.Long | 0.26 | 0.71 | 0.83 | 0.51 | 0.21 |
| PONDR.VLXT | 0.23 | 0.69 | 0.70 | 0.60 | 0.41 |
| PONDR.XL1 | 0.18 | 0.64 | 0.56 | 0.67 | 0.62 |
| PONDR.CAN | 0.13 | 0.60 | 0.51 | 0.65 | 0.34 |

